# *Toxoplasma*-proximal and distal control by GBPs in human macrophages

**DOI:** 10.1101/2021.08.24.457560

**Authors:** Daniel Fisch, Barbara Clough, Rabia Khan, Lyn Healy, Eva-Maria Frickel

## Abstract

Human guanylate-binding proteins (GBPs) are key players of interferon-gamma (IFNγ)-induced cell intrinsic defense mechanisms targeting intracellular pathogens. In this study we combine the well-established *Toxoplasma gondii* infection model with three *in vitro* macrophage culture systems to delineate the contribution of individual GBP family members to control this apicomplexan parasite. Use of high-throughput imaging assays and genome engineering allowed us to define a role for GBP1, 2 and 5 in parasite infection control. While GBP1 performs a pathogen-proximal, parasiticidal and growth-restricting function through accumulation at the parasitophorous vacuole of intracellular *Toxoplasma*, GBP2 and 5 perform a pathogen-distal, growth-restricting role. We further find that mutants of the GTPase or isoprenylation site of GBP1/2/5 affect their normal function in *Toxoplasma* control by leading to mis-localization of the proteins.

## INTRODUCTION

Human cells can defend themselves against pathogens in a process known as cell-intrinsic immunity (MacMicking 2012). Many proteins participating in this are induced by cytokine signaling such as signaling mediated by exposure to the type II interferon-gamma (IFNγ) (Ivashkiv 2018). Amongst IFNγ-induced proteins are several classes of immune GTPases, including the 63 kDa guanylate-binding proteins (GBPs). Humans possess seven *GBP* genes (*GBP1-7*) located in a cluster on chromosome 1 (Olszewski, Gray and Vestal 2006). All GBPs have a similar structure with a N-terminal globular GTPase domain and an elongated C-terminal helical domain (Prakash *et al*. 2000). The GTPase hydrolyses GTP to GDP which induces conformational changes of the proteins (Ghosh *et al*. 2006; Barz, Loschwitz and Strodel 2019; Ince *et al*. 2020). Furthermore, some GBP family members can also hydrolyze GDP to GMP, a unique feature of these proteins (Schwemmle and Staeheli 1994; Praefcke *et al*. 2004; Abdullah, Balakumari and Sau 2010; Wehner and Herrmann 2010). The human GBPs 1, 2 and 5 have a CaaX-box at their C-terminus, which can be modified with an isoprenyl anchor. This lipid tail, together with other sites of the proteins, e.g. a C-terminal polybasic motif R584-586 (Kohler *et al*. 2020), allows for membrane interaction. Moreover, GBPs are known to form dimers and homo-/hetero-oligomers as well as larger protein aggregates (Britzen-Laurent *et al*. 2010; Kravets *et al*. 2016; Ince *et al*. 2017; Wandel *et al*. 2017; Kutsch *et al*. 2020). Some family members are known to target cytosolic and vacuolar bacterial, viral, or protozoal pathogens within cells which leads to their disruption and exposure (Tretina *et al*. 2019). Other functions of GBPs include modulation of apoptosis and pyroptosis, cytokine production, autophagy, radical production, and energy metabolism (Tretina *et al*. 2019). Altogether, they contribute to efficient control of intracellular pathogens.

One common intracellular pathogen of humans is the apicomplexan parasite *Toxoplasma gondii* (Tg), with roughly 30% of humans suffering from non-symptomatic, persistent infection (Pappas, Roussos and Falagas 2009). Tg grows intracellularly once it has infected a human host, forming its own subcellular compartment known as the parasitophorous vacuole (PV) (Sibley 2011). Within the PV, Tg is protected from detection by cytosolic pattern recognition receptors and the innate immune system (Clough and Frickel 2017). While asymptomatic in immune-competent hosts, where Tg transforms into a dormant infection forming tissue cysts in brain and muscle, the parasite can cause the disease known as toxoplasmosis in immunocompromised individuals. Moreover, recurring ocular infections with Tg are a common morbidity in South America, as are complications upon new infection with Tg during pregnancy (Desmonts *et al*. 1985; Daffos *et al*. 1988; Remington *et al*. 2011). Tg infection control in humans critically depends a cell-mediated immune response and on the cytokine IFNγ (Gazzinelli *et al*. 1993, 1994; Hunter *et al*. 1994; Wilson, Matthews and Yap 2008). Tg is therefore a good model pathogen to assess the function of human GBPs.

Macrophages are key cells of the innate immune system. They derive from monocytes infiltrating an inflamed/infected tissue and serve several purposes: macrophages (1) actively phagocytose pathogens and reduce the infectious burden (Rosales and Uribe-Querol 2017), (2) produce cytokines that prime the immune response (Wynn, Chawla and Pollard 2013), (3) present antigens for activation of the adaptive immune response (Roche and Furuta 2015; Hughes *et al*. 2016), (4) clear debris from dead cells (Green, Oguin and Martinez 2016) and (5) contribute to healing of damaged tissues (Feghali and Wright 1997; Cronkite and Strutt 2018). IFNγ which is produced in large amounts during a cell-mediated immune response (Dinarello 2007; Turner *et al*. 2014), activates and polarizes macrophages, and is the key inducer-cytokine for GBPs (Cheng *et al*. 1985; Darnell, Kerr and Stark 1994; Boehm *et al*. 1998). Therefore, GBP-expressing macrophages frequently encounter Tg and are a well-suited model cell line to study GBP functions with sufficient physiological relevance.

Several model cell lines and systems are used to study macrophage biology. One of the most used is the monocytic cancer cell line THP-1 (Chanput, Mes and Wichers 2014). Since long-term culture induces unwanted genetic drift, culture of THP-1 is usually restricted to fewer passages. THP-1 monocytes can be terminally differentiated using phorbol 12-myristate 13-acetate (PMA), a small molecule, irreversible activator of PKC (Ryves *et al*. 1991). Hence, PMA needs to be employed at the minimal concentration necessary for differentiation, in order to reduce activation of cells and so avoid masking any effects of further activations (Park *et al*. 2007). Use of THP-1 cells allows for genome-editing but has the disadvantage of using immortalized cells. Newer systems instead use induced pluripotent stem cells (iPSC). The KOLF iPS cell line can be maintained in culture indefinitely and can be transformed into embryonic bodies (EBs), which upon addition of a cytokine cocktail work as monocyte production factories. Monocytes can be harvested weekly or fortnightly and then terminally differentiated into macrophages with M-CSF (Wilgenburg *et al*. 2013). This produces primary-like human cells. Lastly, primary cells can be used for macrophage biology research. To obtain these, leukocytes are enriched from healthy donor blood. From this, peripheral blood mononuclear cells (PBMCs) can be purified by density-gradient centrifugation from which monocytes are isolated based on surface expression of CD14. These can be terminally differentiated into monocyte-derived macrophages (MDMs). Since MDMs are primary cells, they most accurately reflect human biology. Combining these systems allows for an optimal *in vitro* study of human macrophage biology (Tedesco *et al*. 2018).

In this study, we combine the three distinct macrophage models with gene silencing, genome engineering and high-throughput imaging to delineate the contribution of human GBPs and their mutants to control of Tg infection. We demonstrate that isoprenylated GBPs control the in-part uncoupled processes of Tg growth restriction and parasite killing, critically depending on their correct subcellular localization. Using panels of GBP mutants, we show that GTPase activity and isoprenylation dictate GBP localization and their pathogen-proximal and -distal roles in cell-intrinsic immunity.

## RESULTS

### Human GBP1, 2 and 5 restrict *Toxoplasma* growth in human macrophages and human GBP1 reduces *Toxoplasma* parasite vacuole numbers

To study the function of human GBPs in the context of controlling Tg infection, we utilized three distinct human macrophage culture systems: PMA-differentiated THP-1 macrophages, KOLF iPSCs and *in vitro* differentiated macrophages of purified primary CD14^+^ monocytes from blood of healthy donors (**Figure S1A**). Flow cytometry analysis confirmed presence of the surface markers CD14, FcγRIII (CD16) and CD68 in all macrophage models (**Figure S1B**). RT-qPCR analysis of GBP expression after IFNγ-treatment of the cells showed induction of expression for GBP1 through 5, but no expression of GBP6 or GBP7 in any of the three macrophage models (**Figure S1C+D**) (Fisch *et al*. 2019a). In all cells, GBP1 and GBP2 had the highest total expression levels, followed by GBP5 (**Figures S1C**). Interestingly, the non-isoprenylated GBPs 3 and 4 had the lowest total expression levels in all macrophage models (**Figures S1C**). Of all GBPs, GBP5 showed the highest IFNγ-inducibility, which can be explained by the near complete absence of its transcript in naïve macrophages (**Figure S1D**). GBP3 consistently showed the lowest expression induction (**Figure S1D**).

Having confirmed expression of GBP1 through 5 following IFNγ-treatment of human macrophages, we next used our previously established RNA interference assay, to specifically deplete cells of individual GBP transcripts (Fisch *et al*. 2019a) and assessed their influence on Tg-growth control using high-throughput imaging and analysis with HRMAn (Fisch *et al*. 2019b, 2021). With this assay we could establish that silencing of *GBP1, GBP2* or *GBP5* expression led to a loss of parasite growth restriction (**Figure 1A**) and replication restriction (**Figure 1B**) in all cell lines tested. No GBP appeared to have an influence on the overall proportion of infected cells (**Figure 1C**). Depletion of GBP1 additionally reduced the ability of all IFNγ-primed macrophages to kill intracellular parasites, as measured by determining the vacuole:cell ratio, while GBP2 and 5 contributed to this function to a lesser extent in THP-1 and iPSC macrophages only (**Figure 1D**). Thus, we concluded that GBP1, 2 and 5-depletion significantly restricts Tg growth in 3 macrophage models, while GBP1 additionally kills Tg in all three macrophage models by reducing the vacuole/cell ratio.

**Figure 1:**
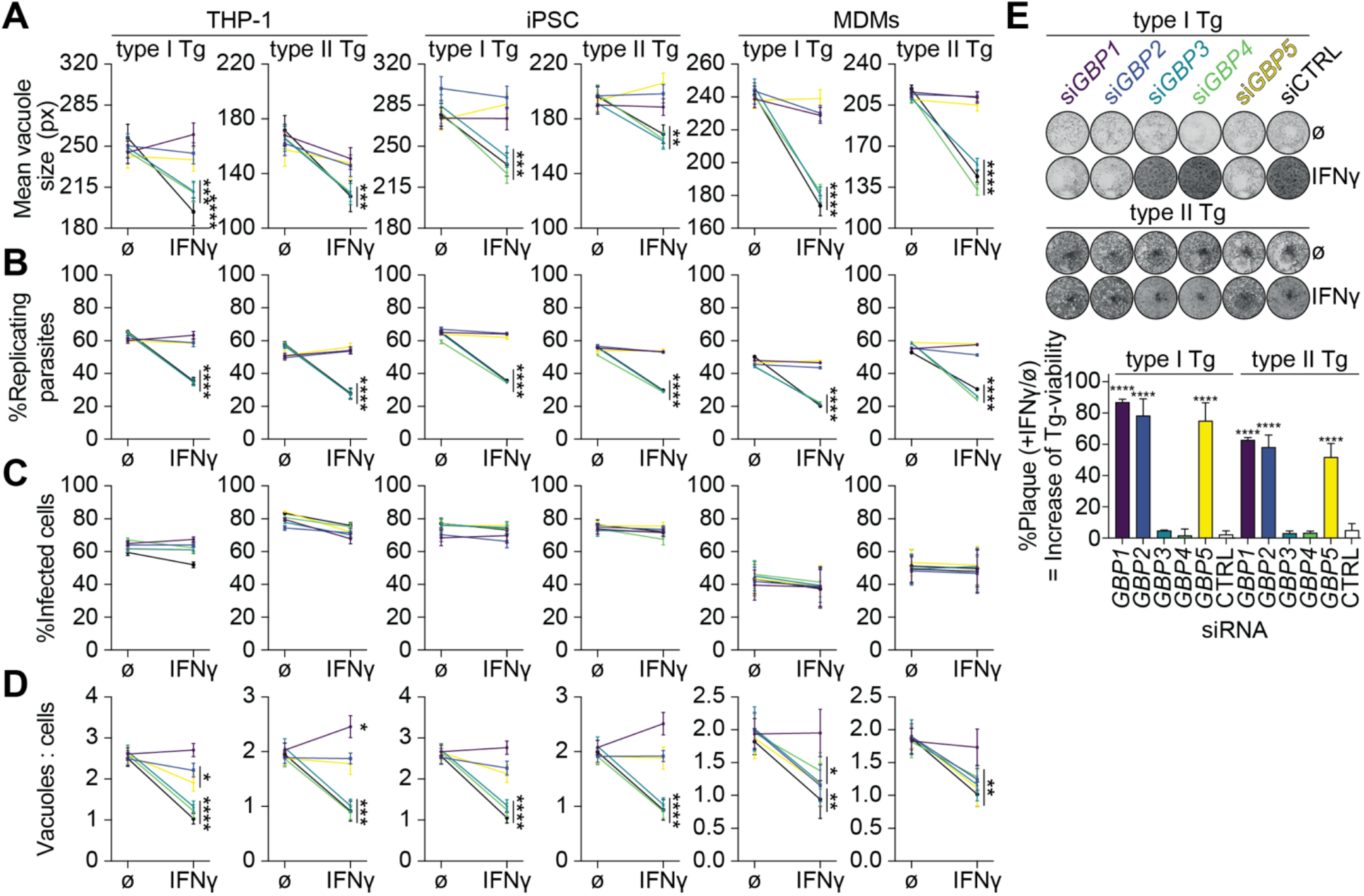
Selective human GBPs limit *Toxoplasma* parasite numbers and restrict their growth in human macrophages. HRMAn-based quantification of mean vacuole size **(A)**, proportion of replicating parasites **(B)**, proportion of infected cells **(C)** and ratio between vacuoles and cells **(D)** of THP-1, induced pluripotent stem cell (iPSC)-derived or monocyte-derived macrophages (MDM) transfected with siRNA against the indicated *GBP* or non-targeting control (CTRL), untreated or primed with IFNγ and infected with type I (RH) or type II (PRU) *Toxoplasma gondii* (Tg) at 18 hours p.i. **(E)** Images of HFF plaques formed by Tg obtained following growth in naïve or IFNγ-primed THP-1 cells for 18 hours additionally transfected with siRNAs against the indicated *GBP* or CTRL (top) and quantification of plaque area from images (bottom). Plaque area of Tg normalized to the corresponding naïve condition, to represent the increase of Tg-viability caused by silencing of the respective *GBP*. **Data information:** Graphs in **(A-D)** shown mean ± SEM from n = 3 independent experiments or n = 4 donors (MDMs). Images in **(E)** representative of n = 3 independent experiments. * *P* ≤0.05; ** *P* ≤ 0.01; *** *P* ≤ 0.001; **** *P* ≤ 0.0001 in **(A-D)** from two-way ANOVA comparing unprimed to IFNγ-primed condition and in **(E)** comparing to CTRL siRNA transfected cells following adjustment for multiple comparisons.

To scrutinize these results obtained using our high-throughput imaging approach, we also determined Tg fitness with traditional plaque assays (**Figure 1E**). We could confirm our observation that GBP1, GBP2 and GBP5 exert Tg-growth control in IFNγ-primed THP-1 macrophages (**Figure 1E**).

### Addition of IFNγ is necessary for restoring *Toxoplasma* growth restriction, but not parasite vacuole numbers when re-expressing GBPs in knockout cells

To further assess the influence of GBP1, 2 and 5 in controlling Tg infection in macrophages, we next assessed THP-1 CRISPR knockout cell lines of the respective gene. THP-1Δ*GBP1* and Δ*GBP5* were previously published (Krapp *et al*. 2016; Fisch *et al*. 2019a) and Δ*GBP2* cells were created using the LentiCRISPR-v2 system. All cell lines were characterized by immunoblotting (**Figure S2A**), RT-qPCR (**Figure S2B**), genotyping PCRs (**Figure S2C**) and Sanger sequencing (**Figure S2D+E**) to confirm absence of the protein and no off-target effects on the other GBP family members. Of note, the knockout cells were created with different approaches, where the *GBP1* gene has a major truncation, *GBP2* is entirely deleted and *GBP5* has nonsense mutations, all rendering the respective gene product absent (**Figure S2**). Next, we used our previously described Doxycycline (Dox)-inducible system (Fisch *et al*. 2019a) and reconstituted the knockout cells with the respective GBP family member (**Figure S2F**). Using these cells in our high-throughput imaging assay, we were able to replicate the previous observation of a loss of Tg-growth and replication restriction in the Δ*GBP1*, Δ*GBP2* and Δ*GBP5* cells which could be reversed by expression induction through addition of Dox (**Figure 2A+B**). Interestingly, addition of Dox alone (expression of just the single GBP) was not sufficient and additional IFNγ-treatment was required (**Figure 2A+B**). This might indicate that several GBPs act in concert or that another IFNγ-inducible factor is required. For parasite killing on the other hand, GBP1 expression alone through Dox-induction could reverse the loss of vacuole/cell control (**Figure 2C**). This effect, however, was only fully restored to wildtype levels upon the extra addition of IFNγ (**Figure 2C**). Complete ablation of GBP2 and 5 by CRISPR in THP-1 macrophages in contrast to downregulation by siRNA showed that these two GBPs are in fact not able to kill Tg via control of the vacuole/cell ratio (**Figure 1D and 2C**). Using HRMAn we further assessed the overall effect of GBP1, 2 and 5 on the total parasite load per cell (**Figure 2D**). This measure combines replication-restriction and killing, and it was only reduced comparable to IFNγ-primed THP-1 WT, if Δ*GBP1*, Δ*GBP2* and Δ*GBP5* cells were treated with IFNγ+Dox. This again shows that for the overall control of the parasite burden GBP1 is essential for killing and growth restriction, whereas GBP2 and GBP5 were needed solely for growth restriction (**Figure 2D**). In summary, human macrophages express GBPs 1 through 5 upon IFNγ-stimulation and GBP1, 2 and 5 all contribute to the growth control of the intracellular parasites, while GBP1 is additionally responsible for controlling vacuole/cell numbers.

**Figure 2:**
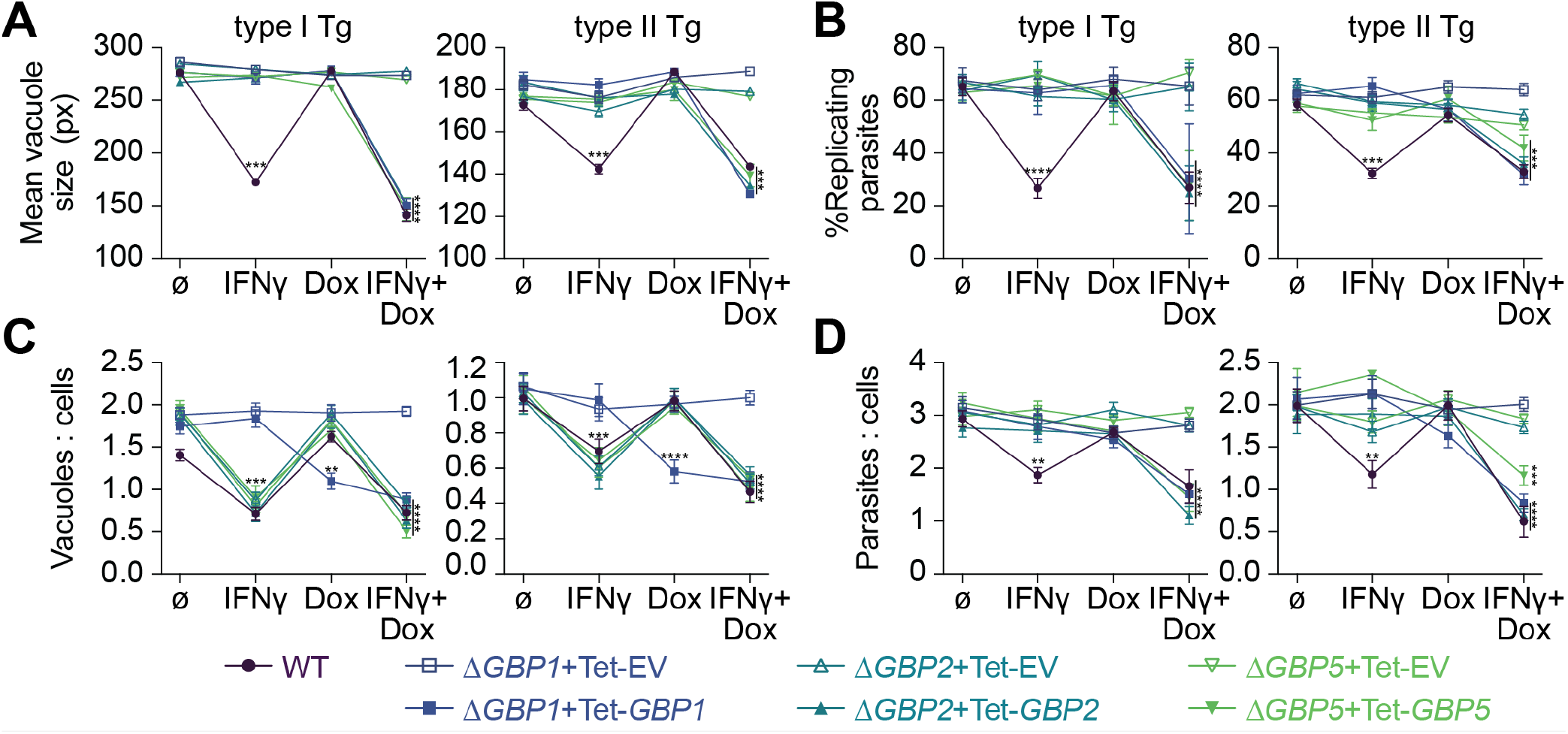
Re-expression of GBPs restores *Toxoplasma* restriction in Δ*GBP* cells. HRMAn-based quantification of mean vacuole size **(A)**, proportion of replicating parasites **(B)**, ratio between vacuoles and cells **(C)** and ratio between parasites and cells **(D)** of THP-1Δ*GBP1*, Δ*GBP2* or Δ*GBP5* cells transduced with Tet-empty vector (EV, open symbols) or Tet-*GBP1/2/5* (closed symbols) untreated or primed with IFNγ and/or Doxycycline (Dox) and infected with type I (RH) or type II (PRU) *Toxoplasma gondii* (Tg) at 18 hours p.i. **Data information:** Graphs in **(A-D)** shown mean ± SEM from n = 3 independent experiments. * *P* ≤ 0.05; ** *P* ≤ 0.01; *** *P* ≤ 0.001; **** *P* ≤ 0.0001 for indicated condition in **(A-D)** from two-way ANOVA comparing to untreated condition following adjustment for multiple comparisons.

### GTPase activity and lipidation of GBP1, 2 and 5 are essential for their anti-*Toxoplasma* activity

We next created panels of mutants for GBP1, GBP2 and GBP5 targeting their GTPase activity, C-terminal lipidation, the polybasic motif in GBP1 and its dimerization capacity (**Figure 3A**). We transduced the respective Δ*GBPx* cells with the Dox-inducible system (**Figure S3**). We then assessed the effect of these mutants on the functionality of the proteins (**Figure 3B-D**). To do so, we performed our high-throughput imaging assay as before by treating THP-1 macrophages with IFNγ+Dox and normalized the resulting effects to the IFNγ-only treated control of the same cell line. In this way, the only difference is presence or absence of the wildtype or mutated GBP protein in otherwise IFNγ-primed cells. Like this, we were able to calculate the proportion of Tg-growth restriction or killing of the respective GBP functionality relative to the absence of the same GBP (**Figure 3B-D**).

**Figure 3:**
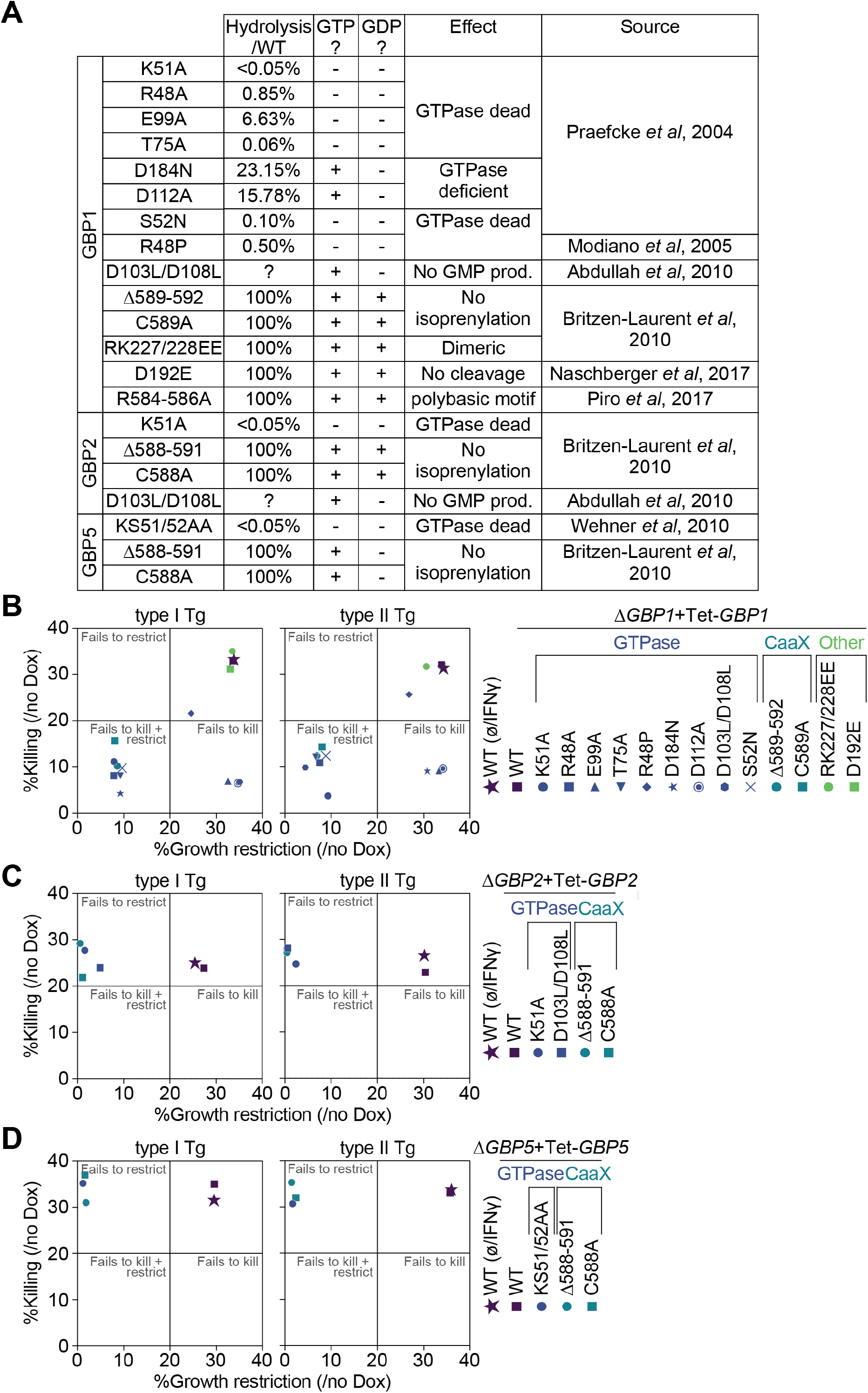
GTPase activity and lipidation of GBP1, 2 and 5 are essential for their anti-*Toxoplasma* activity. Overview of GBP1, GBP2 and GBP5 mutants **(A)**. Growth restriction and killing (= ratio between vacuoles and cells) of type I (RH) and type II (PRU) *Toxoplasma gondii* (Tg) at 18 hours p.i. in THP-1Δ*GBP1*+Tet-*GBP1* cells expressing the indicated mutant of GBP1 **(B)**, Δ*GBP2*+Tet-*GBP2* cells expressing the indicated mutant of GBP2 **(C)**, Δ*GBP5*+Tet-*GBP5* cells expressing the indicated mutant of GBP5 **(D)** or of IFNγ-treated THP-1 WT cells for each, plotted as proportion between IFNγ + Doxycycline (Dox)-treated versus IFNγ-only-treated cells. **Data information:** Graphs in **(B-D)** show mean from n = 3 independent experiments.

Screening the GBP1 mutants showed that mutations rendering the GTPase activity non-functional (K51A, R48A, T75A, D184N or S52N) failed to restrict Tg growth and killing, whereas GTPase-mutants that predominantly affected GMP-production (E99A, D112A or D103L/D108L) still restricted the growth but failed to kill Tg. GBP1^R48P^ with a predicted inactive GTPase was still active to restrict and kill Tg, although slightly impaired in this capacity (**Figure 3B**). Isoprenylation site mutations (C589A or Δ589-592) also failed to kill and restrict Tg-growth (**Figure 3B**).

GBP2 and GBP5 mutations that abolish GTP hydrolysis (K51A or D103L/D108L for GBP2 and KS51/52AA for GBP5) or mutations of the isoprenylation sites (C588A or Δ588-591 for both) failed to restrict Tg-growth (**Figure 3C+D**). Since neither protein contributes to Tg-killing, this was unaffected and likely carried out by endogenous GBP1 induced through IFNγ-priming of the cells (**Figure 3C+D**).

### GBP2 and 5 do not localize to *Toxoplasma* vacuoles

Comparing findings of the GBP mutant screen indicates a close link between GBP1 GTPase activity/isoprenylation and the control of Tg reminiscent of previous results on GBP1 recruitment and correlation to Tg-killing and host cell death (Fisch *et al*. 2019a, 2020). This suggests a functional link between these processes. Thus, mCherry-tagged GBPx mutants, showing a pathogen growth control phenotype, were created, and transduced into Δ*GBP*x+Tet cells to study the localization and spatiotemporal activities (**Figure S4**). Using mCH-GBP1 WT, mCH-GBP2 WT and mCH-GBP5 WT expressing cells, we could confirm that GBP5 was localizing to the Golgi apparatus as had been described before in epithelial cells (Tripal *et al*. 2007; Britzen-Laurent *et al*. 2010) (**Figure 4A**). In IFNγ-primed, uninfected cells, GBP1 mutants of the GTPase or isoprenylation site appeared more dispersed in the cytosol instead of showing a granular appearance like GBP1 WT. This might indicate a loss of membrane interactions or aggregate formation (**Figure 4B**). The GBP1^R584-586A^ mutant of the polybasic motif had the most dramatic effect with the protein forming large aggregates within uninfected cells (**Figure 4B**) as has been observed before (Kohler *et al*. 2020; Kutsch *et al*. 2020). The observed dispersed cytoplasmic localization of GBP2 had no obvious differences with mutation of the protein, but GBP5 mutants affecting the GTPase or its isoprenylation had lost their localization at the Golgi (**Figure 4B**).

**Figure 4:**
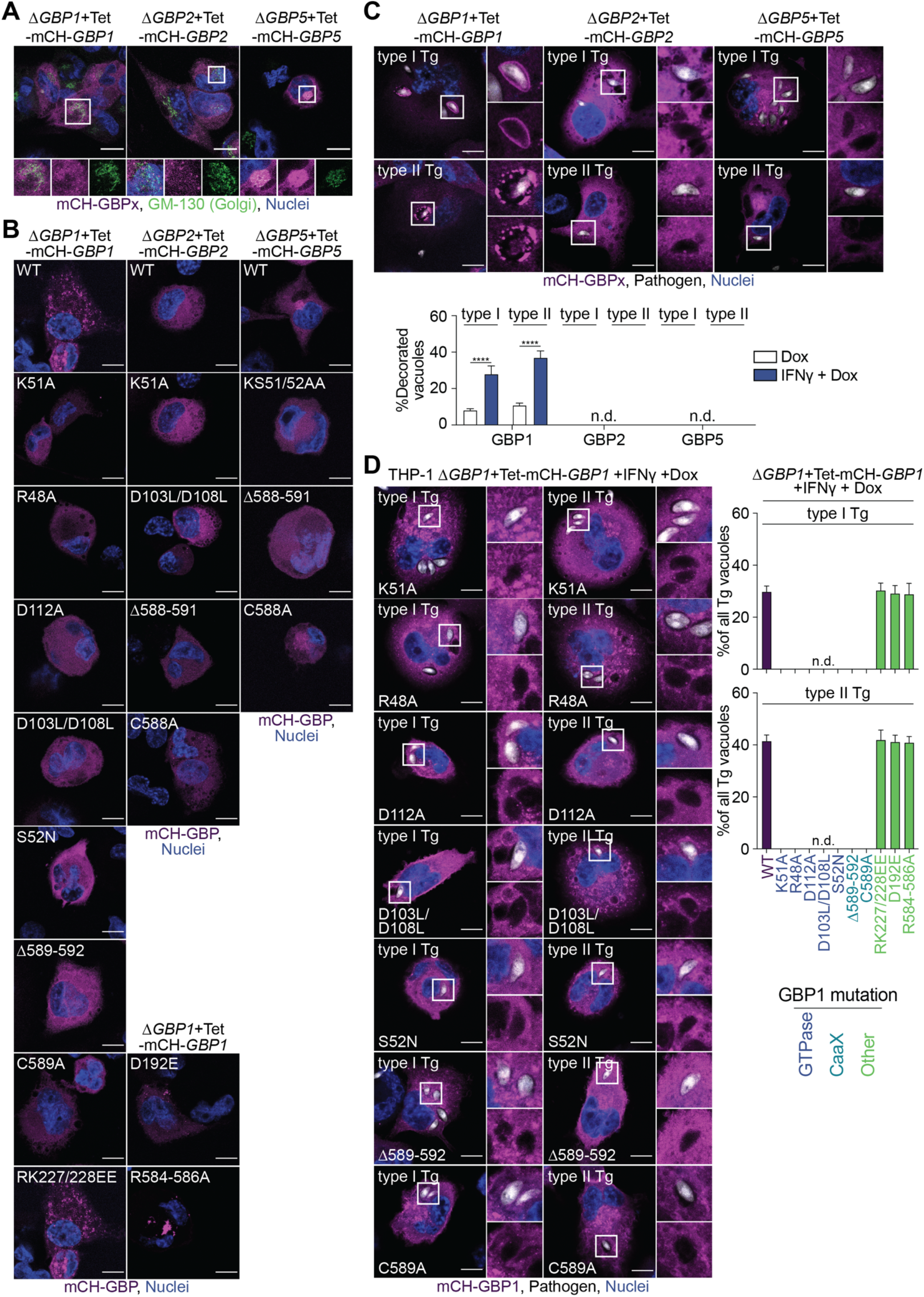
GBP2 and 5 do not localize to *Toxoplasma* vacuoles. **(A)** Immunofluorescence images of THP-1Δ*GBP1*+Tet-mCH-*GBP1*, Δ*GBP2*+Tet-mCH-*GBP2* or Δ*GBP5*+Tet-mCH-*GBP5* cells treated with IFNγ and Doxycycline (Dox) and stained for Golgi marker GM-130 to illustrate Golgi localization of GBP5 in uninfected cells. Magenta: mCherry (mCH)-GBP1/2/5; Green: GM-130 (Golgi); Blue: nuclei. Scale bar, 10 μm. **(B)** Immunofluorescence images of THP-1Δ*GBP1*+Tet-mCH-*GBP1*, Δ*GBP2*+Tet-mCH-*GBP2* or Δ*GBP5*+Tet-mCH-*GBP5* cells expressing the indicated GBPx mutant treated with IFNγ+Dox to illustrate localization of the respective protein in uninfected cells. Magenta: mCherry (mCH)-GBP1; Blue: nuclei. Scale bar, 10 μm. **(C)** Immunofluorescence images (top) and HRMAn-based quantification of GBP recruitment to Tg (bottom) in THP-1Δ*GBP1*+Tet-mCH-*GBP1*, Δ*GBP2*+Tet-mCH-*GBP2* or Δ*GBP5*+Tet-mCH-*GBP5* cells treated with IFNγ+Dox and infected with type I (RH) or type II (PRU) *Toxoplasma gondii* (Tg) for 6 hours. Magenta: mCherry (mCH)-GBP1/2/5; Grey: Tg; Blue: nuclei. Scale bar, 10 μm. **(D)** Images (left) and HRMAn-based quantification of GBP1 recruitment to Tg-vacuoles (right) in THP-1Δ*GBP1*+Tet-mCH-*GBP1* cells expressing the indicated GBP1 mutant treated with IFNγ+Dox and infected for 6 hours. Magenta: mCherry (mCH)-GBP1; Grey: pathogen; Blue: nuclei. Scale bar, 10 μm. **Data information:** Images in **(A+C-D)** representative of n = 3 and in **(B)** representative of n = 2 independent experiments. Graph in **(C+D)** show mean ± SEM from n = 3 independent experiments. **** *P* ≤ 0.0001 for indicated comparisons in **(C)** from one-way ANOVA comparing to Dox-only treated cells following adjustment for multiple comparisons; n.d. not detected.

In contrast to GBP1, neither GBP2 WT nor GBP5 WT recruited to Tg vacuoles in infected human macrophages implying that they have their growth restrictive function away from the pathogen (pathogen-distal) (**Figure 4C**). We therefore only assessed recruitment of GBP1 mutants to Tg. In agreement with our previous observations of correlation between modulation of macrophage cell death and GBP1 recruitment to pathogens (Fisch *et al*. 2019a, 2020), all GBP1 GTPase and isoprenylation mutants failed to target Tg vacuoles in IFNγ-primed THP-1 cells (**Figure 4D**).

In summary, GBP1, GBP2 and GBP5 contributed to the control of Tg infection via parasite growth restriction and reduction of vacuole/cell numbers in three different human, *in vitro* macrophage models, including primary-like iPSCs and primary MDMs. Genome engineering and use of a Dox-inducible system confirmed GBP1 targeting to pathogen vacuoles to depend on its GTPase activity and isoprenylation. Other infection- and IFNγ-treatment-dependent factors are probably involved in regulating its Tg control function as well. Furthermore, GBP1 needs to be able to produce GMP and be targeted to vacuoles to kill Tg parasites by reducing vacuole/cell numbers. Surprisingly, GBP2 and GBP5 did not target Tg vacuoles, but were involved in Tg growth restriction. This function depended on both GBP2 and 5 GTPase activity and isoprenylation.

## DISCUSSION

Here, we employed three *in vitro* models to study the role of human GBPs in infected macrophages. Gene depletion experiments in THP-1 cells, MDMs and iPSC macrophages established that GBP1, GBP2 and GBP5 control the replication of Tg, while GBP1 was additionally parasiticidal. The findings on pathogen control by GBPs were confirmed using THP-1 CRISPR KO cell lines and rescued by reconstituting protein expression. Use of an imaging-based assay also allowed to delineate the contribution of individual GBPs to restriction and/or killing, and extend observations made by overall pathogen burden assessment through classical plaque formation assays.

Following IFNγ-stimulation, macrophages express GBP1 through 5, but not GBP6 or GBP7, which are predominantly expressed in the oropharyngeal tract (Uhlen *et al*. 2015) and which was expected since GBP6/7 lack GAS elements in their promoter regions (Tretina *et al*. 2019). GBP expression patterns resembled expression profiles in mesenchymal stem cells (Qin *et al*. 2017). Our findings furthermore concur with previous studies showing an effect of human GBP1 on Tg growth in mesenchymal stem cells and in A549 lung epithelial cells (Johnston *et al*. 2016; Qin *et al*. 2017). A role for human GBP2 and GBP5 in Tg infection control has so far not been established, but a large body of literature suggests and supports a similar role for their murine homologues (Virreira Winter *et al*. 2011; Kravets *et al*. 2012, 2016; Degrandi *et al*. 2013; Matta *et al*. 2018).

The three GBP family members that can be isoprenylated (Nantais *et al*. 1996; Tripal *et al*. 2007; Britzen-Laurent *et al*. 2010) contributed to Tg growth restriction, while GBP3 and GBP4 did not. Moreover, these three GBPs were highly upregulated and expressed upon IFNγ-stimulation, while GBP3 and GBP4 show significantly lower expression and inducibility in all three human macrophage models studied here. This may indicate a different role for GBP3/4. One conceivable hypothesis is that GBP3/4 regulate lipidated GBPs through heterotypic interactions, partially resembling the Irg system of the mouse, in which GMS-Irgs control the activity of the GKS-Irgs (Hunn *et al*. 2008; Haldar *et al*. 2013, 2016).

It is likely that GBP1, 2 and 5 act in concert. siRNA-depletion and Dox-reconstitution experiments suggest that for growth restriction all three GBPs are needed, since depletion of a single member abolished restriction and conversely reconstitution of a single member did not rescue the loss of restriction in the CRISPR KO cells. Growth restriction alone was not able to reduce the overall parasite burden. For this to occur, Tg-killing mediated by GBP1 was required. Similar hierarchical organization of the human GBP system was observed during *Shigella flexneri* infection where the pathfinder GBP1 first targets the pathogen thus facilitating recruitment of GBP2/3 and GBP4 (Piro *et al*. 2017; Wandel *et al*. 2017). It is likely that similar cooperation is needed between GBP1, 2 and GBP5 for their pathogen-distal action against Tg. Additionally, it is probable that GBP1 also has a pathogen-distal function for Tg growth restriction, as mutants that cannot produce GMP do not localize to the PV but still restrict the parasite growth. These GBP1 mutants therefore resemble the function of GBP2/5.

In uninfected cells GBP1, GBP2 and GBP5 showed differing localizations: GBP1 had a granular appearance suggesting aggregate formation or (endo-)membrane interaction, GBP2 was uniformly distributed in the cytosol and GBP5 associated with the Golgi apparatus, resembling prior observations in HeLa cells (Britzen-Laurent *et al*. 2010). It is known that correct localization of the three isoprenylated GBPs depends on lipidation with farnesyl (GBP1) or geranylgeranyl (GBP2/5) (Britzen-Laurent *et al*. 2010). Accordingly, mutation of the CaaX box of either of the three GBPs led to uniform cytoplasmic distribution. Since GBP1/2/5 all show differing subcellular localizations despite all being isoprenylated, other parts of the proteins must contribute to their correct trafficking. One example could be the polybasic motif of GBP1 (R584-586), which when mutated led to the pronounced phenotype of protein aggregation, as observed by other groups too (Kohler *et al*. 2020; Kutsch *et al*. 2020).

GBP1, 2 and 5 have all been localized at the Golgi in previous studies (Modiano, Lu and Cresswell 2005; Tripal *et al*. 2007; Britzen-Laurent *et al*. 2010; Krapp *et al*. 2016; Braun *et al*. 2019). Aluminium fluoride treated HeLa cells or HFFs showed accumulation of GBP1 at the Golgi, suggesting that this only occurs in a GTP-locked conformation (Modiano, Lu and Cresswell 2005). GBP5 has a well-established localization at the Golgi and can further recruit GBP2 (Britzen-Laurent *et al*. 2010; Braun *et al*. 2019). In line with our results, isoprenylation of GBP5 was required for this. Localization of GBP5 at the Golgi is needed for its antiviral activity against HIV (Krapp *et al*. 2016), which is achieved by concerted action of GBP2 and GBP5, together reducing the activity of Furin protease (Braun *et al*. 2019). Since GBP5 GTPase and isoprenylation mutants lost their association with the Golgi apparatus, it is likely that GBP5 activity against Tg relies on its correct localization to the Golgi. Thus, GBP1/2/5 influence Tg growth by acting without accumulation of the proteins at the pathogen (pathogen-distal), which has been observed before for GBP1 in A549 lung-epithelial cells (Johnston *et al*. 2016) but contests the dogma of defense protein accumulation at the intracellular infection site (MacMicking 2012). It is tempting to speculate that the GBPs therefore have additional functions during infection other than recruiting to pathogens.

Apart from Tg-restriction mechanism(s), GBP1 accumulated at Tg vacuoles in infected cells. Neither GBP2 nor GBP5 recruited to Tg. The recruitment of GBP1 was dependent on its GTPase function and isoprenylation. GBP1 recruitment might also rely on other proteins, as its association with Tg appears cell-type- and IFNγ-dependent. It will therefore be interesting to study GBP1-interactomes. Comparative study of macrophage and A549 lung epithelial cell GBP1-interactomes might offer the opportunity to identify critical GBP1 trafficking factors. Overall, recruitment of GBP1 to Tg resembles the function of its murine homologue, which is known to associate with bacterial pathogens (Kim *et al*. 2011; Haldar *et al*. 2014; Meunier *et al*. 2014, 2015; Finethy *et al*. 2015; Man *et al*. 2015; Feeley *et al*. 2017; Wallet *et al*. 2017; Zwack *et al*. 2017; Lindenberg *et al*. 2017; Balakrishnan *et al*. 2018; Liu *et al*. 2018) and Tg-PVs and was also found directly on the parasites (Virreira Winter *et al*. 2011; Kravets *et al*. 2012, 2016; Degrandi *et al*. 2013; Haldar *et al*. 2014, 2015; Costa Franco *et al*. 2018).

Careful examination of the effect of different mutations of the GBP1 GTPase activity (Praefcke *et al*. 2004; Modiano, Lu and Cresswell 2005; Abdullah, Balakumari and Sau 2010) revealed that full GTPase activity was needed for recruitment to Tg and killing of the pathogen, while GMP formation was dispensable for growth restriction. Interestingly, GBP1 was the only parasiticidal GBP family member, a function which may therefore rely on the formation of GMP. Similar observations have been made for *Chlamydia* infections, where GMP formation was necessary for pathogen and host-cell killing, but dispensable for *Chlamydia* growth restriction (Xavier *et al*. 2020). Conversely, GBP5 which cannot produce GMP by hydrolysis of GDP (Wehner and Herrmann 2010), did not kill Tg. GBP2 however, which like GBP1, can hydrolyze GDP to GMP (Abdullah, Balakumari and Sau 2010), did not kill Tg. GTPase activity of GBP2 and GBP5 were nevertheless needed for Tg-growth restriction.

Taken together these results show that killing of Tg relies on GBP1 recruitment to the pathogens and a pathogen-proximal function involving the formation of GMP, whereas GBP1, 2 and 5 restrict Tg-growth via a thus far unknown pathogen-distal function.

## AUTHOR CONTRIBUTION

DF and EMF conceived the project. DF conducted the experiments. BC set up and assisted with imaging experiments. RK and LH set up initial iPS cell culture. DF and EMF wrote the manuscript. EMF supervised the project. EMF acquired funding related to the project.

## ACKNOWLEDGEMTS

The authors would like to acknowledge all members of the Frickel lab (University of Birmingham, UK) for critical discussion of this study and Mike Howell from the HTS STP at the Francis Crick Institute for his help with high-throughput image acquisition.

## MATERIALS AND METHODS

### Cell and parasite culture, treatments, and infection

THP-1 (TIB202, ATCC) were maintained in RPMI with GlutaMAX (35050061, Gibco) and 10% FBS (Sigma), HFFs (SCRC 1041, ATCC) and HEK293T (Cell Services, The Francis Crick Institute, London, UK) were maintained in DMEM with GlutaMAX and 10% FBS at 37°C in 5% CO_2_. THP-1s were differentiated with 50 ng mL^-1^ phorbol 12-myristate 13-acetate (PMA, P1585, Sigma) for 3 days and then rested for 2 days in PMA-free, complete medium. All cells were regularly tested for mycoplasma by immunofluorescence and PCR. Cells were stimulated for 16 h prior to infection with addition of 50 IU mL^-1^ human IFNγ (285-IF, R&D Systems). Induction of GBP expression in the Dox-inducible cells was performed with 200 ng mL^-1^ Dox overnight (D9891, Sigma).

Tg were maintained by serial passage on HFF cell monolayers and passaged onto new HFFs the day before infection. Tg were prepared from freshly 25G syringe lysed cultures by centrifugation at 50 x g for 3 minutes, transferring the cleared supernatant into a new tube, subsequent centrifugation at 500 x g for 7 minutes and re-suspension of the pelleted parasites into fresh complete medium. Parasite-suspension was added to the cells at a MOI of 1. The cell cultures with added Tg were then centrifuged at 500 x g for 5 minutes to synchronize infection. Two hours post-infection, extracellular parasites were removed with three PBS washes (806552, Sigma) and fresh complete medium added prior to culturing at 37°C, 5% CO_2_ for the required time.

### iPS cell culture and monocyte/macrophage production

Production of monocytes from KOLF iPSC (HESCU STP, The Francis Crick Institute, London, UK) was previously described (Wilgenburg *et al*. 2013). KOLF cells were maintained in their pluripotent state in a feeder-free, serum-free culture system at 37°C in 5% CO_2_ using Synthemax™ II-SC Substrate-coated plates (3535, Corning) and mTeSRTM-1 medium (85850, StemCell Technologies). Cells were clump-passaged when colonies covered ∼75% of the wells by washing with PBS, detaching using Collagenase IV (07427, StemCell Technologies), followed by gentle scraping in mTeSRTM-1 medium. Cells were split roughly 1:4 and supplemented with 1 mM Rock-inhibitor (Y27632; Calbiochem). Cells were fed with new media daily.

To create monocyte production factories KOLF cells were washed with PBS and harvested with TrypLE Express (12604021, Gibco), dissociated into single cells by pipetting and finally diluted 1:10 with PBS and collected in a centrifuge tube. Cells were pelleted by centrifugation and resuspended in mTeSRTM-1 supplemented with 1 mM Rock-inhibitor, 50 ng mL^-1^ BMP-4 (120-05, Peprotech), 20 ng mL^-1^ SCF (130-093-991, Miltenyi), 50 ng mL^-1^ VEGF (100-20, Peprotech) (= EB medium). Next, AggreWell™ 800 plates (34811, StemCell Technologies) were prepared by rinsing with PBS, addition of 1 mL EB medium to each well and centrifugation at 3,000 x g for 2 minutes. Then 1 mL of harvested cells were added per well, the plate centrifuged at 150 x g for 3 minutes and left in the incubator for four days. EBs were fed daily with fresh EB medium by stepwise exchanging 75% of medium. EBs were harvested by dislodging through pipetting, transferring the well-contents onto a 40 mm strainer, rinsing with PBS and collecting them into a new tube. 500 EBs were transferred per T175 tissue culture flasks in 20 mL X-VIVO™15 (04-418Q, Lonza), supplemented with 100 ng mL^-1^ M-CSF (PHC9504, Gibco), 25 ng mL^-1^ IL-3 (203-GMP, R&D Systems), 2 mM GlutaMAX, 100 U mL^-1^ penicillin/streptomycin (15140122, Invitrogen), and 0.05 mM β-Mercaptoethanol (21985023, Gibco). Roughly 2–3 weeks following seeding, monocytes in suspension appeared and were harvested fortnightly from the supernatant. Monocytes were differentiated into macrophages in X-VIVO™15 supplemented with 100 ng mL^-1^ M-CSF for 5 days.

### Primary human macrophage isolation and culture

PBMCs were extracted from Leukocyte cones from healthy donors (NHS) via Ficoll (17544202, GE Healthcare) density gradient centrifugation. CD14^+^ monocytes were extracted using magnetic microbeads (130-050-201, MACS Miltenyi). Monocytes were counted, seeded, and differentiated for one week in RPMI containing 10% human AB serum (H4522, Sigma), GlutaMAX, penicillin/streptomycin and 5 ng mL^-1^ hGM-CSF (130-093-864, Miltenyi). The medium was replaced after 2 and 5 days, to replenish the hGM-CSF.

### siRNA transfection

Cells were transfected two days prior to infection, at the time the THP-1 differentiation medium was replaced, or MDM/iPSC differentiation medium was replaced on day 5 after seeding. All siRNAs were used at a final concentration of 30 nM. To set up the transfection mix, a 10x mix was prepared in OptiMEM containing the appropriate siRNA(s) and TransIT-X2 transfection reagent (MIR 600x, Mirus) in a 1:2 stoichiometry. As the GBPs exhibit high sequence similarity, a costume transfection panel using three different Silencer™ Select siRNAs (Ambion: GBP1: s5620, s5621, s5622; GBP2: s5623, s5624, s5625; GBP3: s5626, s5627, s5628; GBP4: s41805, s41806, s41807; GBP5: s41808, s41809, s41810) was used (Fisch *et al*. 2019a). The appropriate negative control was Silencer™ Select Negative Control No. 1 siRNA (#4390843, Ambion).

### Plaque assays

0.8×10^6^ differentiated THP-1 cells were infected with Tg as described above and 18 hours p.i. supernatant and cells were harvested from the wells of a 12-well plate. Cells were syringe-lysed and obtained parasites from within the cells and the supernatant diluted 1:10,000 and added to HFFs grown confluent in wells of a 24-well plate.

Determination of plaque sizes and number was performed 5 days p.i. of the HFFs, when cells were fixed with ice-cold methanol and stained with crystal violet (C6158, Sigma). Following 5 washes with PBS, plaques were imaged on a GelCount™ Colony Counter (Oxford Optronix) and cell covered area determined using FIJI. Proportions of plaque and plaque loss, as compared to Tg grown in untreated THP-1, were calculated.

### Flow cytometry

1×10^6^ differentiated macrophages were harvested using accutase (A6964, Sigma) and scraping and washed twice with warm PBS. Cells were resuspended in PBS + 1% BSA containing dilutions of fluorescently labelled antibodies against surface receptors and incubated for 1 hour at room temperature in the dark. Cells were washed with PBS, fixed with 4% formaldehyde for 15 minutes at room temperature and washed again, prior to resuspension in PBS + 1% BSA. All samples were analyzed on a LSR Fortessa (BD Biosciences), and recorded data was processed using FlowJo 10.3 (FlowJo, LLC).

### RT-qPCR

RNA was extracted from 0.25×10^6^ cells using Trizol reagent (15596026, Invitrogen). 5 μg mL^-1^ GlycoBlue (AM9516, Invitrogen) was added during the isopropanol (190764, Sigma) precipitation to increase RNA-yields. RNA quality was measured on a Nanodrop 2000 Spectrophotometer (Thermo Scientific). 1 μg RNA was reverse transcribed using high-capacity cDNA synthesis kit (4368813, Applied Biosystems). qPCR used PowerUP SYBR green (A25742, Applied Biosystems), 20 ng cDNA in a 20 μL reaction and primers at 1 μM final concentration on a QuantStudio 12K Flex Real-Time PCR System (Applied Biosystems). Primer specificity was ensured by designing primers to span exon-exon junctions, whenever possible, and for each primer pair a melt curve was recorded (see **Table 2**). Ct values were normalized to the Ct of human *HPRT1*. To determine absolute expression of GBPs, a defined amount of linearised plasmid standards was added as PCR template and obtained Ct values used to calculate transcript numbers from the samples.

**Table 1:**
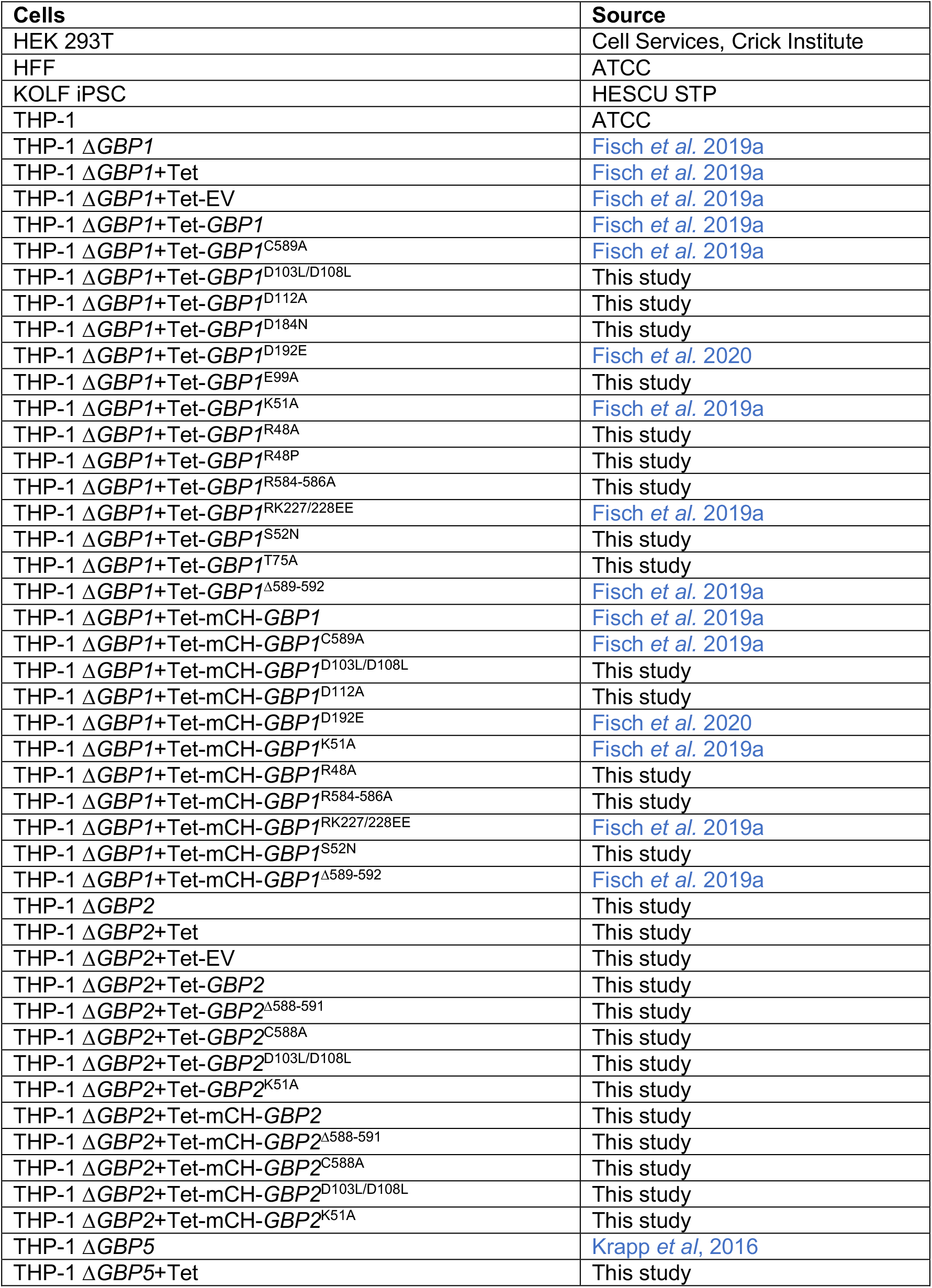
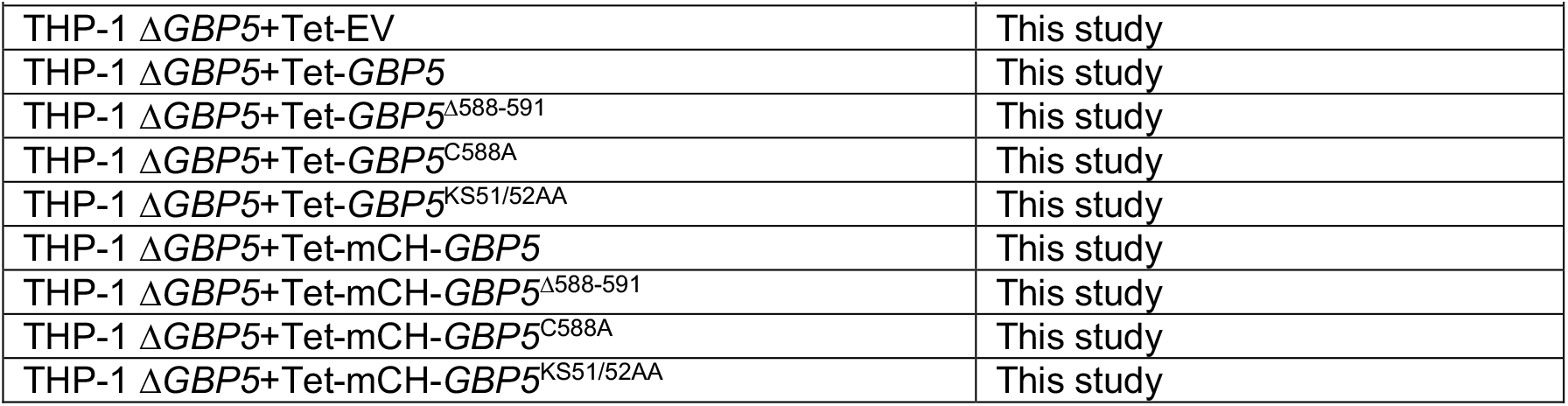
List of cell lines.

**Table 2:**
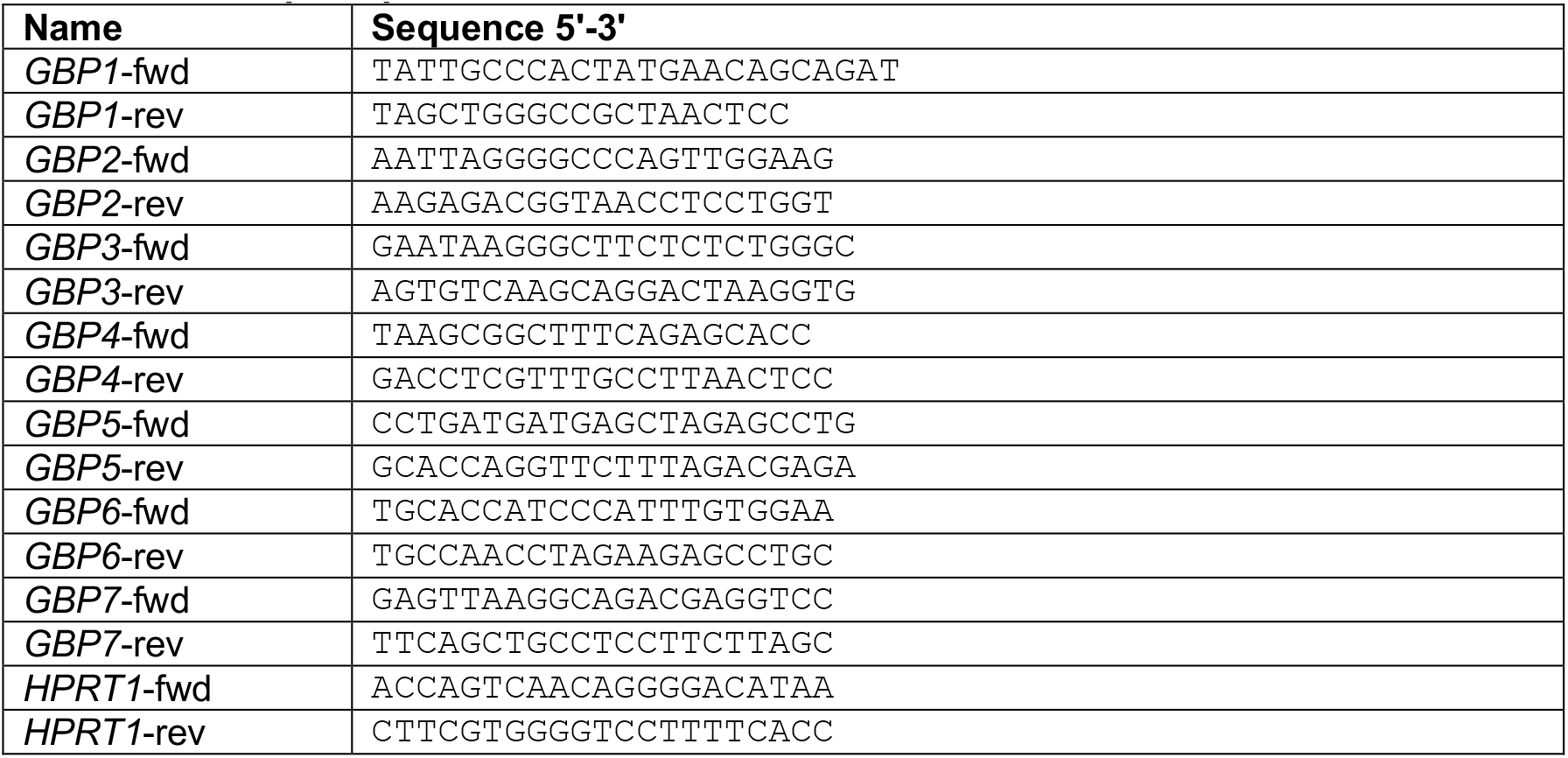
List of qPCR primers.

### Creation of new cell lines

THP-1Δ*GBP1* and the Dox-inducible system were previously published (Fisch *et al*. 2019a). THP-1Δ*GBP5* were a gift from Frank Kirchhoff (Krapp *et al*. 2016). Guide RNA (gRNA) sequences targeting the 5’ and 3’ UTR of *GBP2* gene were designed using cripr.mit.edu. DNA oligonucleotides encoding for the crRNAs (sgRNA1: 5’-CACCGTGTCTTACAAATTGGG TCAC-3’; sgRNA2: 5’-CACCGCATGAGTTGAATTGC TCTGT-3’) were annealed by mixing in equimolar ratios and boiling at 95°C for 15 minutes followed by a slow decrease to room temperature. Annealed oligos were then cloned into BsmBI-digested (ER0451, Thermo Scientific) pLentiCRISPR-V2 backbone (Sanjana, Shalem and Zhang 2014) using Quick Ligation™ kit (M2200, NEB) and transduced into THP-1 WT cells using Lentiviral particles (Fisch *et al*. 2019a). Following selection with 1 μg mL^-1^ Puromycin (A1113802, Gibco) for 14 days, cells were sub-cloned by serial dilution into ten 96-well plates using pre-conditioned complete medium supplemented with non-essential amino acids (11140076, Gibco), penicillin/streptomycin and GlutaMAX. Roughly 3 weeks after seeding of the single cells, obtained clones were expanded into 24-well plates with 2 mL fresh medium and screened for absence of *GBP2* expression by RT-qPCR. Clones that showed reduced or absent *GBP2* expression underwent secondary screening by immunoblotting. Finally, confirmed KO clones were tested again by Sanger sequencing of the genomic target locus, RT-qPCR, and immunoblotting.

Cells with Dox-inducible expression were created as previously published (Fisch *et al*. 2019a). To create plasmids that express GBP1, GBP2 or GBP5 under the control of Dox, RNA from IFNγ-treated THP-1s was extracted and cDNA synthesized as described above. The CDS of GBP mRNA was amplified with Q5 polymerase, the amplicons treated with Taq polymerase (M0273, NEB) to create A-overhangs and cloned into pCR2.1^®^-TOPO TA vector using TOPO TA kit (451641, Invitrogen). GBP mutants were created by site-directed mutagenesis, introducing single point mutations with mismatch-primers and PCR with Q5 polymerase. Using the mutated or wildtype GBP-containing vectors, the ORFs were PCR-amplified to create overhangs to pLenti-Tet vector. Gibson assemblies of the digested backbone and the GBP ORFs were performed, and successful cloning confirmed by Sanger sequencing. To create mCH-tagged versions, mCH-ORF was amplified with overlaps to the backbone and the GBP ORF and included in the Gibson assembly reactions. GBP ORFs lacking the C-terminal CaaX-box were amplified with primers excluding parts of the wildtype GBP ORFs.

### SDS-PAGE and immunoblotting

0.5×10^6^ cells were seeded per well of a 48-well plate, differentiated, and treated as described above. At the end of treatments, cells were washed with ice-cold PBS and lysed for 5 min on ice in 50 μL RIPA buffer (150 mM NaCl, 1% Nonidet P-40, 0.5% Sodium deoxycholate, 0.1% SDS, 25 mM Tris-HCl pH 7.4) supplemented with protease inhibitors (Protease Inhibitor Cocktail set III, EDTA free, Merck) and PhosSTOP phosphatase inhibitors (4906845001, Roche). Lysates were cleared by centrifugation at full speed for 15 minutes at 4°C. BCA assay (Pierce BCA protein assay kit, 23225, Thermo Scientific) was performed to determine protein concentrations. 10 μg of total protein per sample were mixed with Laemmli buffer (#1610737, Biorad) containing 5% DTT (646563-10X, Sigma) and boiled at 95°C for 10 minutes and then run on Bis-Tris gels (Novex, Invitrogen) in MOPS running buffer.

Following SDS-PAGE, proteins were transferred onto Nitrocellulose membranes using iBlot transfer system (Invitrogen). Membranes were blocked with either 5% BSA (A2058, Sigma) or 5% dry-milk (M7409, Sigma) in TBS-T (0.05% Tween-20) for at least 1 h at room temperature. Incubation with primary Abs (see **Table 3**) was performed at 4°C overnight. Blots were developed by washing the membranes with TBS-T, probed with 1:5,000 diluted HRP-conjugated secondary Abs in 5% BSA in TBS-T and washed again. Finally, the membranes were incubated for 2 minutes with ECL (Immobilon Western, WBKLS0500, Millipore) and chemiluminescence recorded on a ChemiDoc MP imaging system (Biorad).

**Table 3:**
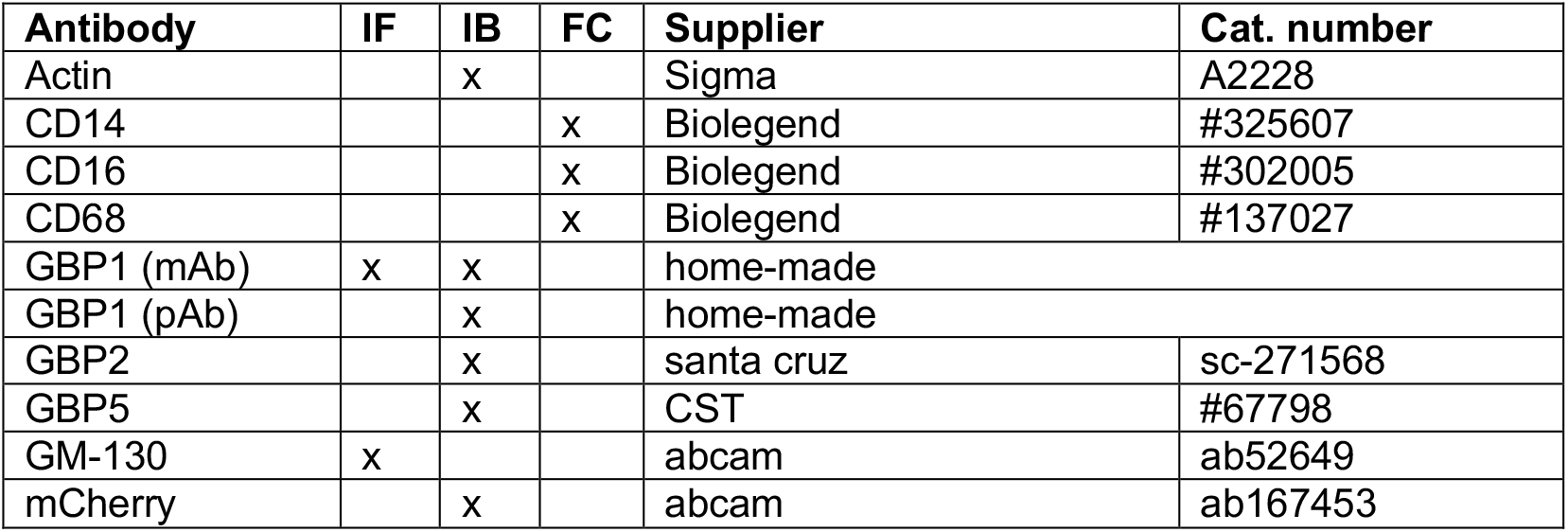
List of primary antibodies. IF: immunofluorescence; IB = immunoblotting; FC: flow cytometry

### Microscopy

0.25×10^6^ cells were seeded on gelatin-coated (G1890, Sigma) coverslips in 24-well plates. Following differentiation, treatments and infection, cells were washed three times with warm PBS, prior to fixation, to remove any uninvaded pathogens and then fixed with 4% methanol-free formaldehyde (28906, Thermo Scientific) for 15 min at room temperature. For high-throughput imaging 50,000 cells were seeded of a black-wall, clear bottom 96-well imaging plate (Thermo Scientific), differentiated, and treated and fixed as described above.

Following fixation, cells were washed again with PBS and kept at 4°C overnight to quench any unreacted formaldehyde. Fixed specimens were permeabilized with PermQuench buffer (0.2% (w/v) BSA and 0.02% (w/v) saponin in PBS) for 30 minutes at room temperature and then stained with primary Abs (see **Table 3**) for one hour at room temperature. After three washes with PBS, cells were incubated with the appropriated fluorescently labelled secondary Ab and 1 μg mL^-1^ Hoechst 33342 (H3570, Invitrogen) diluted in PermQuench buffer for 1 hour at room temperature. Cells were washed with PBS five times and mounted using 5 μL Mowiol. For high-throughput imaging, fixed and permeabilized specimens were stained for 1 hour at room temperature by adding PermQuench buffer containing 1 μg/mL Hoechst 33342 and 2 μg mL^-1^ CellMask™ Deep Red plasma membrane stain (H32721, Invitrogen). After staining, the specimens were washed with PBS five times and kept in 200 μL PBS per well for imaging.

Coverslips were imaged on a Leica SP5-inverted confocal microscope using 100x magnification and analyzed using LAS-AF software. Plates were imaged on a Cell Insight CX7 High-Content Screening (HCS) Platform (Thermo Scientific) using 20x magnification. Following acquisition, images were exported from HCS Studio Cell Analysis as single channel 16-bit .tiff files before they were fed into the HRMAn analysis pipeline (Fisch *et al*. 2019b, 2021).

### Data handling and statistics

Data was plotted using Prism 8.4.0 (GraphPad Inc.) and presented as means of n = 3 experiments (with usually 3 technical repeats within each experiment) with error bars as standard error of the mean (SEM), unless stated otherwise. Significance of results was determined by non-parametric one-way ANOVA or two-way ANOVA as indicated in the figure legends. Benjamini, Krieger and Yekutieli false-discovery rate (Q = 5 %) based correction for multiple comparisons as implemented in Prism was used when making more than 3 comparisons.

## Supplementary Information

This is supplementary information to the preprint manuscript from Fisch *et al*. 2021. Supplementary section contains supplementary tables, supplementary figures and figure legends. All further information is available upon request.

### Supplementary tables

### Supplementary Figures

**Figure S1:**
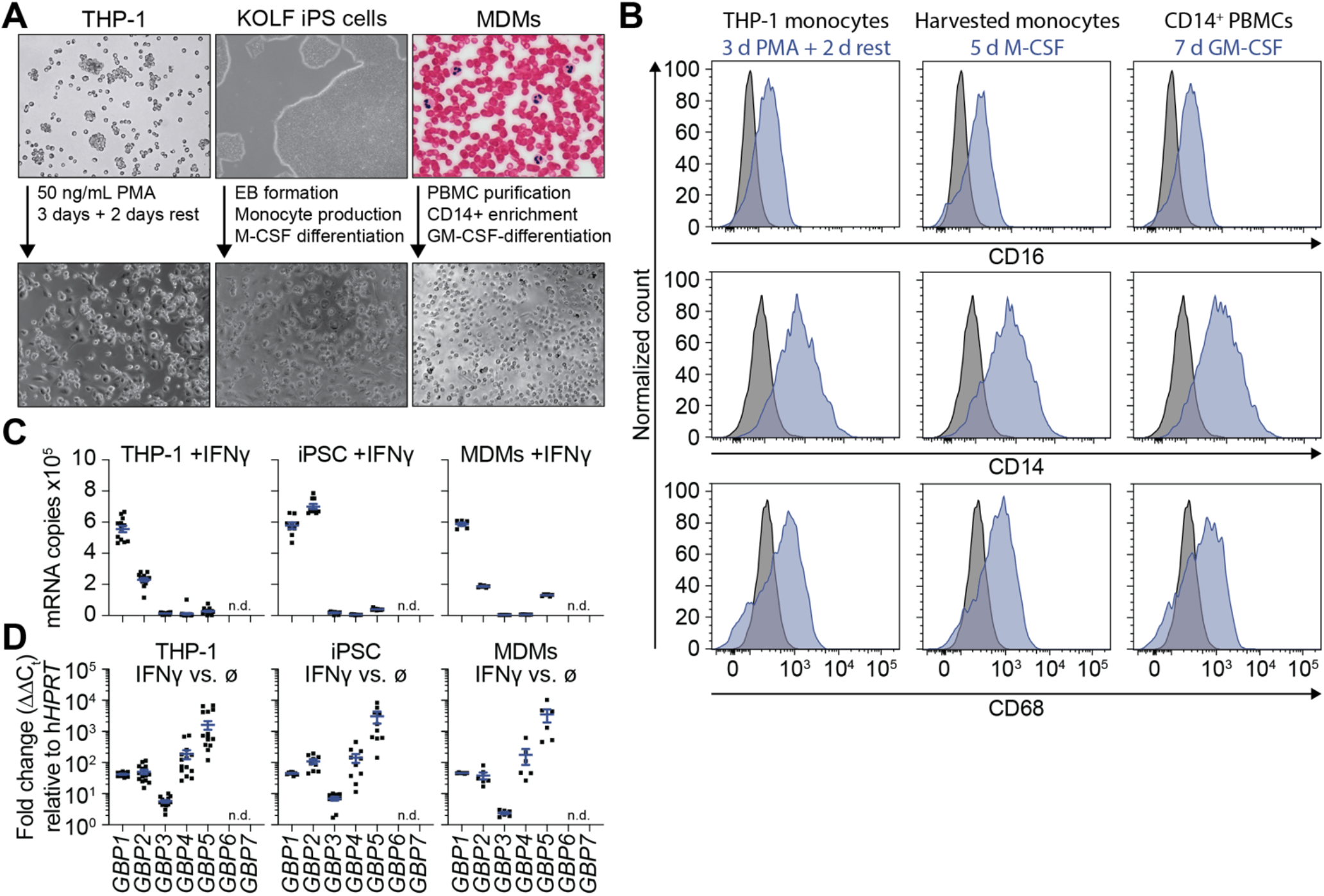
Growth morphology and differentiation of human macrophage models. **(A)** Images of human macrophage cell lines before (top) and following indicated differentiation steps (bottom). Human induced pluripotent stem cells (iPSC); embryonic body (EB); peripheral blood monocytic cell (PBMC). Scale bar, 50 μm. Image of whole blood from https://commons.wikimedia.org/. **(B)** Flow cytometry measuring CD14, CD16 or CD68 surface expression of the indicated cell line, prior to (black) or post differentiation into macrophages (blue). **(C)** Absolute mRNA copies of indicated *GBP* per cell for IFNγ-primed THP-1, iPSC-derived or monocyte-derived macrophages (MDM) at 16 h post treatment. **(D)** Fold change induction of indicated *GBP* following treatment of indicated cell line with 50 IU mL^-1^ IFNγ for 16 hours, plotted as ΔΔCt normalizing to untreated cells (ø) and relative to human Hypoxanthine-guanine phosphoribosyltransferase (*HPRT1*). **Data information:** Graphs in **(B)** representative of n = 3 independent experiments or n = 4 different donors (MDMs). Graphs in **(C+D)** show data from n = 14 (THP-1) or n = 10 (iPSC) independent experiments or n = 6 donors (MDMs) and mean ± SEM; n.d. not detected.

**Figure S2:**
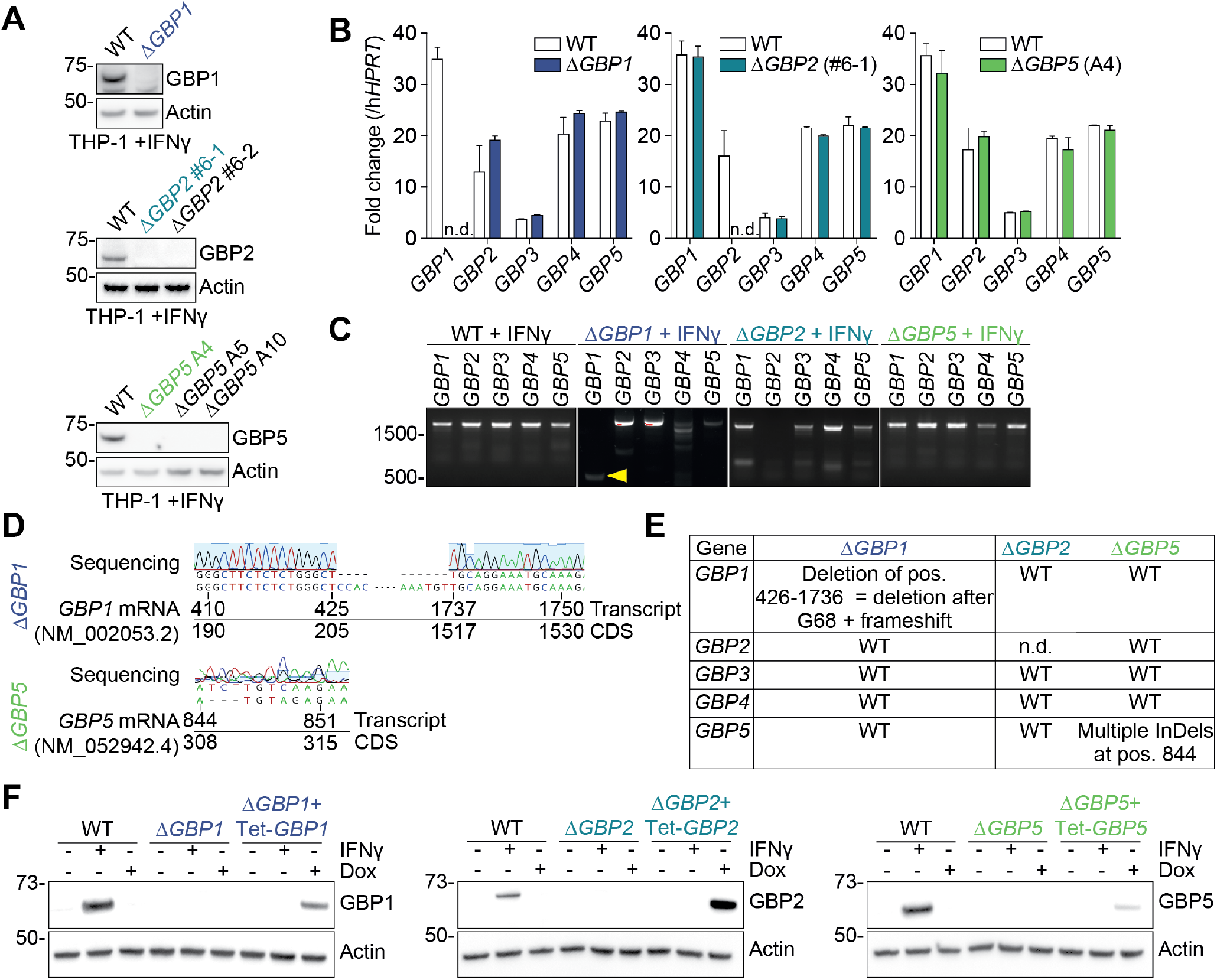
Characterization of human *GBP* CRISPR knockout cell lines. **(A)** Immunoblots for indicated proteins from IFNγ-primed THP-1 WT, THP-1Δ*GBP1*, Δ*GBP2* or Δ*GBP5* cells. **(B)** Fold change of mRNA expression of indicated human *GBP* in IFNγ-primed THP-1 WT, THP-1Δ*GBP1*, Δ*GBP2* or Δ*GBP5* cells relative to human Hypoxanthine-guanine phosphoribosyltransferase (*HPRT1*); n.d.: not detected. **(C)** PCR amplification of indicated *GBP* coding sequences (CDS) on cDNA created from mRNA obtained from IFNγ-primed THP-1 WT, THP-1Δ*GBP1*, Δ*GBP2* or Δ*GBP5* cells. Yellow arrowhead highlights truncated GBP1 CDS in THP-1Δ*GBP1* cells. **(D)** Sanger sequencing of *GBP1* and *GBP5* CDS captured into TOPO-TA vectors following PCR amplification shown in **(C)** aligned to displayed wildtype sequence of respective *GBP* mRNA. **(E)** Overview of sequencing results of human *GBP* CDS from THP-1Δ*GBP1*, Δ*GBP2* or Δ*GBP5* cells; n.d. not detected. **(F)** Immunoblots from THP-1 WT, Δ*GBP1*, Δ*GBP1*+Tet-*GBP1*, Δ*GBP2*, Δ*GBP2*+Tet-*GBP2*, Δ*GBP5* and Δ*GBP5*+Tet-*GBP5* cells treated with IFNγ, and Doxycycline (Dox) as indicated. **Data information**: Images in **(A), (C)** and **(D)** representative of n = 2 experiments. Graphs in **(B)** show mean ± SEM from n = 3 independent experiments. Images in **(F)** representative of n = 3 independent experiments.

**Figure S3:**
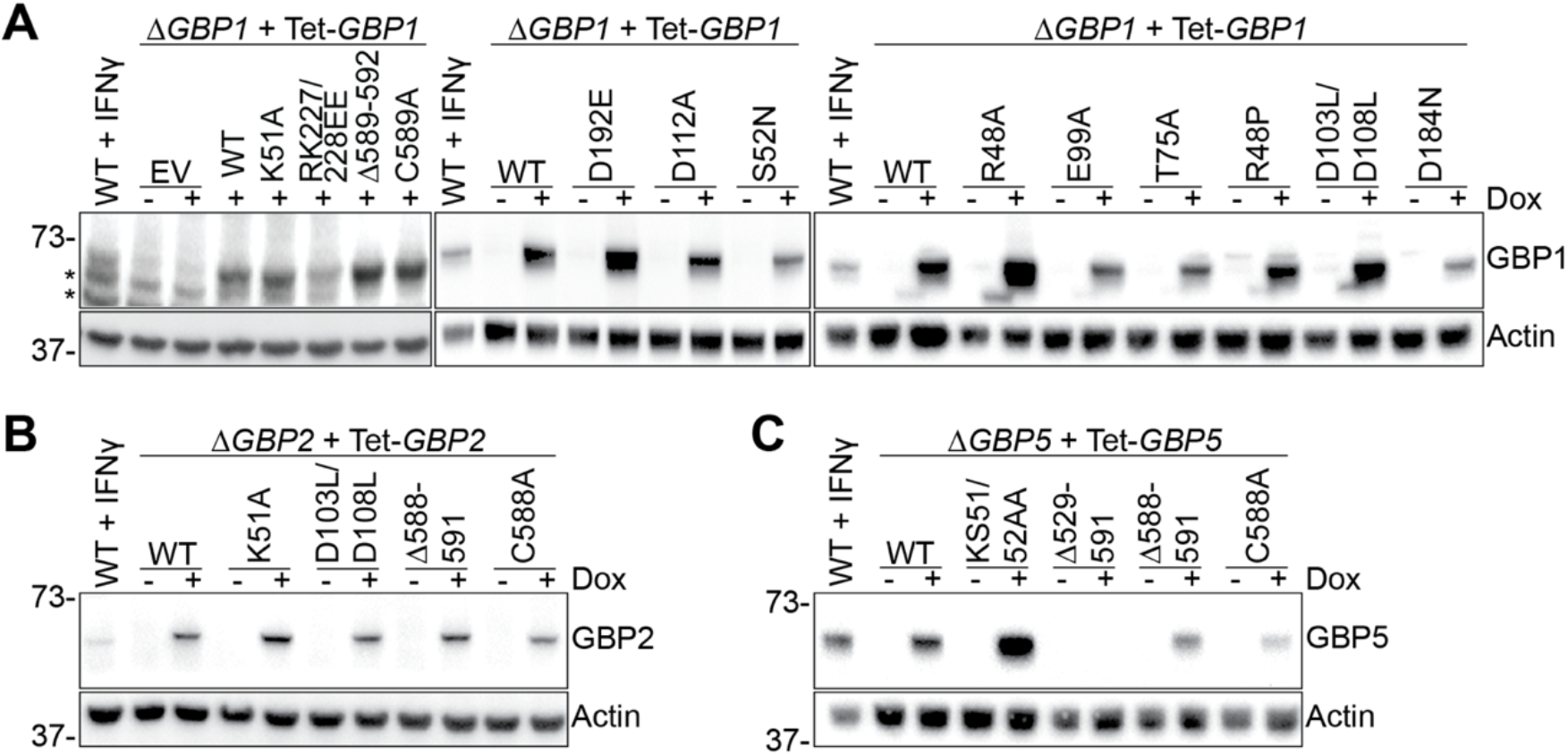
Reconstitution of mutated GBPs into macrophages. Immunoblots from IFNγ-primed THP-1 WT and Δ*GBP1* **(A)**, Δ*GBP2* **(B)** or Δ*GBP5* cells **(C)** reconstituted with the indicated mutants of GBP1, GBP2 or GBP5 respectively and treated with Doxycycline (Dox) as indicated. *Marks unspecific bands.

**Figure S4:**
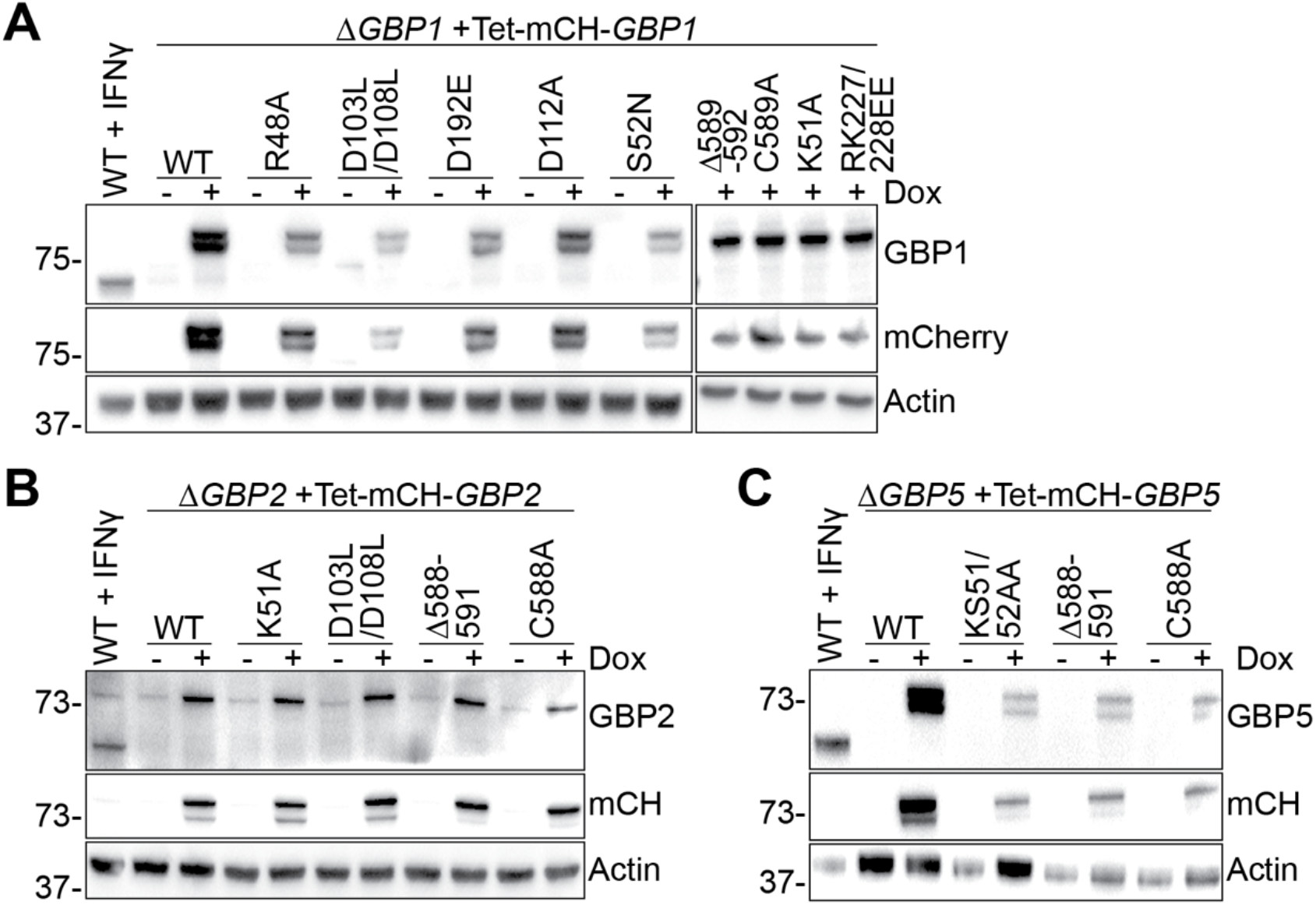
Reconstitution of mutated mCherry-tagged GBPs into macrophages. Immunoblots from IFNγ-primed THP-1 WT and Δ*GBP1*, Δ*GBP2* or Δ*GBP5* cells reconstituted with the indicated mCherry (mCH)-tagged mutants of GBP1, GBP2 or GBP5 respectively and treated with Doxycycline (Dox) as indicated.

